# Minimizing the number of optimizations for efficient community dynamic flux balance analysis

**DOI:** 10.1101/2020.03.12.988592

**Authors:** James D. Brunner, Nicholas Chia

**Author notes:** Mayo Clinic, 200 First St. SW, Rochester, MN, USA.

## Abstract

Dynamic flux balance analysis uses a quasi-steady state assumption to calculate an organism’s metabolic activity at each time-step of a dynamic simulation, using the well-know technique of flux balance analysis. For microbial communities, this calculation is especially costly and involves solving a linear constrained optimization problem for each member of the community at each time step. However, this is unnecessary and inefficient, as prior solutions can be used to inform future time steps. Here, we show that a basis for the space of internal fluxes can be chosen for each microbe in a community and this basis can be used to simulate forward by solving a relatively inexpensive system of linear equations at most time steps, instead of the full optimization problem. Using our method, we can use this solution as long as the resulting metabolic activity remains within the optimization problem’s constraints (i.e. the solution remains feasible). As the solution becomes infeasible, it first becomes a feasible but degenerate solution to the optimization problem, and we can solve a different but related optimization problem to choose an appropriate basis to continue forward simulation. We show using an eight species community that this is an efficient and robust method for computing dynamic flux balance analysis simulations, and so is capable of simulating communities of organisms. We demonstrate that the method gives an approximately 85% speed-up per organism over the standard and widely used method. Our method has been implemented in the *Python* language and source code is available at https://github.com/jdbrunner/surfin_fba and in the Python Package Index (PyPI) as `surfinFBA`.

**Author summary:** The standard method in the field for dynamic flux balance analysis carries a prohibitively high computational cost because it requires solving a linear optimization problem at each time-step. We have developed a novel method for producing solutions to this dynamical system which greatly reduces the number of optimization problems that must be solved. We prove mathematically that we can solve the optimization problem once and simulate the system forward as an ordinary differential equation for some time interval, and solutions to this ODE provide solutions to the optimization problem. Eventually, the system reaches an easily checkable condition which implies that another optimization problem must be solved. We compare our method with the classical method to validate that it provides equivalent solutions in much lower computational time.

## Introduction

### Microbial communities and human health

The makeup of microbial communities is often complex, dynamic, and hard to predict. However, microbial community structure has a profound effect on human health and disease [1–7]. These two facts have lead to significant interest in mathematical models which can predict relative abundances among microbes in a community. Recently, advances in genetic sequencing have allowed the creation of genome-scale models (GEMs) which reflect the internal network of cellular metabolism, and can therefore be used to predict metabolite use and production [8–10]. This technique can be extended to microbial community modeling by combining GEMs of different species. There has been significant interest in using GEMs to predict relative populations of stable microbial communities [11–17]. Community metabolic modeling can not only predict relative populations, but also holds the potential to predict and explain the community metabolite yield, which can have a profound effect on health [4]. Furthermore, model repositories such as the online bacterial bioinformatics resource *PATRIC* [18] or the *BiGG model database* [19] make it possible to build community models using information from individual species investigations.

Additionally, various dynamical models have been proposed to explain and predict microbial community population dynamics [20–24]. Among these are models which propose that interactions between species are mediated by the metabolites that each species produces and consumes [25, 26], and there is significant evidence that these models perform better than models which depend on direct interaction between species [27, 28].

GEMs can be used to predict microbial growth rates as well as metabolite consumption and production rates using a process called *flux balance analysis* (FBA). Because these predictions appear in the form of rates of change, they can be used to define a metabolite mediated dynamical model, simply by taking as a vector field the rates of change predicted by FBA. We can therefore combine the techniques of metabolite mediated dynamic modeling and community metabolic modeling to produce dynamic predictions of microbial community population size and metabolite yield. This strategy is called *dynamic FBA* [29, 30], and has recently been used to model microbial communities [31, 32].

Dynamic FBA, when implemented naively, requires a linear optimization problem to be repeatedly solved, and so carries a high computational cost even for small communities. Furthermore, *in silico* experiments may need to be repeated many times over various environmental conditions or using various parameter choices in order to make robust conclusions or to fit accurate model parameters. As a result, a naive implementation of dynamic FBA carries a prohibitively high computational cost to make conclusions about larger microbial communities. The implementation of dynamic FBA in the popular COBRA toolbox software package [8] is done in this way, and so carries a high computational burden. Dynamic FBA can be improved by taking advantage of the linear structure of the optimization problem, so that optimizations are not necessary for every time step [33, 34].

In order to implement dynamic FBA without optimizing at each time step, we use an optimal basic set for the FBA linear optimization problem to create a system of linear equations whose solutions at future time-steps coincide with the solutions to the FBA optimization problem. It is possible that the basic index set chosen may not provide future solutions if the optimization problem’s solution is degenerate. This is potentially a meaningful limitation of the method, because most re-optimizations required by the method involve degenerate solutions. To solve this problem, we provide a method to choose among possible basic index sets for a degenerate solution to the optimization problem. We show that there exists a choice of basic index set that allows forward simulation. We then describe an algorithm that chooses this useable basis in order to simulate forward with as few full optimizations as possible. Additionally, our implementation makes use of a slightly less costly optimization problem. Our method is available as a Python package published in the Python Package Index (PyPI) as `surfinFBA`, and source code is available at https://github.com/jdbrunner/surfin_fba.

In this manuscript, we detail how dynamic FBA can be simulated forward without re-optimization for some time interval, and give a method for doing so. We propose conditions on an optimal basic set for the FBA linear optimization problem which allows for forward simulation, and we prove that such a choice exists. We then detail how to choose this basis set, and finally give examples of simulations which demonstrate the power of our method and its speed relative to the classical implementation introduced in [30].

## Background

### Flux balance analysis

With the advent of genetic sequence and the resulting genome scale reconstruction of metabolic pathways, methods have been developed to analyze and draw insight from such large scale models [9]. To enable computation of relevant model outcomes, constraint based reconstruction and analysis (COBRA) is used to model steady state fluxes *υ*_*i*_ through a microorganism’s internal metabolic reactions under physically relevant constraints [9]. One of the most basic COBRA methods, called *flux balance analysis* (FBA) optimizes some combination of reaction fluxes ∑*γ_i_υ_i_* which correspond to increased cellular biomass, subject to the constraint that the cell’s internal metabolism is at equilibrium:

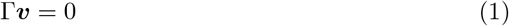

where Γ is the *stoichiometric matrix*, a matrix describing the stoichiometry of the metabolic model.

This optimization is of interest because it reflects the optimization carried out by nature through evolution [9]. The vector *γ* = (*γ*_1_, *γ*_2_, …, *γ*_*d*_) is an encoding of cellular objectives, reflecting the belief that the cell will be optimized to carry out these objectives. The constraint Eq. (1) means that any optimal set of fluxes found by FBA corresponds to a steady state of the classical model of chemical reaction networks [35]. This reflects the assumption that the cell will approach an internal chemical equilibrium.

The optimization is done over a polytope of feasible solutions defined by the inequalities *υ*_*i,min*_ ≤ *υ*_*i*_ ≤ *υ*_*i,max*_, or possibly more complicated linear constraints. See Fig. 2 for a geometric representation of an example of the type of linear optimization problem that is carried out. By convention, forward and reverse reactions are not separated and so negative flux is allowed. Linear optimization problems like FBA often give rise to an infinite set of optimal flux vectors ***υ*** = (*υ*_1_, *υ*_2_, …, *υ*_*d*_). Geometrically, this set will correspond to some face of the polytope of feasible solutions. To draw conclusions despite this limitation, many methods have been developed to either characterize the set of optimal solutions, as with flux variability analysis (FVA), or enforce more constraints on the network to reduce the size of this set, as with loopless FVA [9].

### Dynamic FBA

Flux balance analysis provides a rate of increase of biomass which can be interpreted as a growth rate for a cell. Furthermore, a subset of the reactions of a GEM represent metabolite exchange between the cell and its environment. By interpreting constraints on nutrient exchange reactions within the metabolic network as functions of the available external metabolites and fluxes of exchange reactions as metabolite exchange rates between the cell and its environment, the coupled system can be modeled. The simplest way to do this is to use an Euler method, as in [30]. The algorithm from [30] is summarized in Fig. 1.

**Fig 1.**
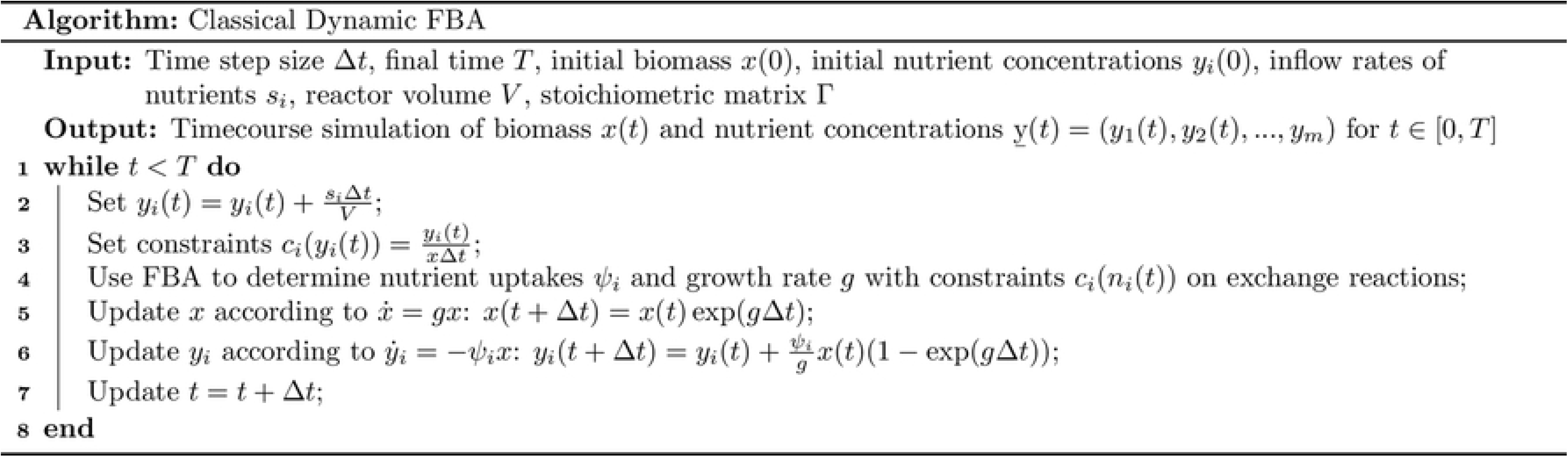
Dynamic FBA algorithm given in [30]. This is an Euler method which simply evolves the community forward with constant growth rates and metabolite exchange rates for a single time-step. The method then must recalculate a solution to the linear optimization problem.

This method is implemented in the COBRA toolbox [8], but notably requires a complete recalculation of the network fluxes *at each timestep*. Furthermore, for complex systems, time-steps must be small to avoid numerical error which causes the simulation to fail entirely. In our simulations of an 8-species system, a time-step of 10^−4^ was necessary to avoid negative metabolite biomasses, which cause the simulation to behave extremely inaccurate if allowed to continue forward. Thus, to simulate to 1 hour of growth, 10^4^ linear programs must be solved for each species.

However, resolving the system at each time step is not necessary, as the solution the optimization problem at some initial time can actually be used to compute future optimal solutions. Höffner et al, [34], used this observation to introduce a variable step-size method for dynamic FBA. In that method a basic index set is chosen by adding biological constraints to the optimization problem hierarchically until a unique optimal solution is found. The challenge of such an approach is in choosing the basis for the optimal solution, as the optimal basis is not guaranteed to be unique. We describe how to choose a basis from among the possibilities provided from an FBA solution which is most likely to remain optimal as simulation proceeds forward. We therefore prioritize reducing the number of times the linear program must be solved.

Additionally, a method described as “dynamic optimization approach” was introduced in Mahadevan et al., [29], however this method is computationally expensive. In particular, the method given in [29] involves optimizing over the entire time-course simulated, and so is formulated as a non-linear program which only needs to be solved once. While this method requires only one optimization, this optimization is itself prohibitively difficult due to the dimensionality of the problem growing with the fineness of time-discretization.

### The dynamic FBA model for communities

We can write a metabolite mediated model for the population dynamics of a community of organisms ***x*** = (*x*_1_, …, *x*_*p*_) on a medium composed of nutrients ***y*** = (*y*_1_, …, *y*_*m*_):

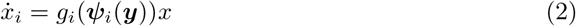

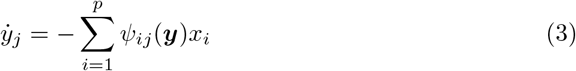

where ***ψ***_*i*_ is a vector of the fluxes of nutrient exchange reactions for organism *x*_*i*_ as determined by FBA. Using FBA to determine ***ψ***_*i*_ is therefore a quasi-steady state assumption on the internal metabolism of the organisms *x*_*i*_ [36–38].

Recall that the basic assumption of flux balance analysis is that, given a matrix Γ_*i*_ that gives the stoichiometry of the network of reactions in a cell of organism *x*_*i*_ that growth *g*_*i*_(***y***) is the maximum determined by solving the following linear program [9]:

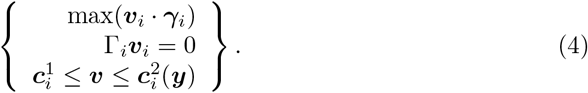

The key observation allowing dynamic FBA is that the optimal solution to this problem also determines ***ψ***_*i*_ simply by taking *ψ*_*ij*_ to be the value of the flux *υ*_*ij*_ of the appropriate metabolite exchange reaction. For clarity, we will relabel the elements of ***υ***_*i*_ so that *ψ*_*ik*_ = *υ_ij_* if *υ*_*ij*_ is the *k*^*th*^ exchange flux, and *ϕ*_*ik*_ = *υ*_*ij*_ if *υ*_*ij*_ is the *k*^*th*^ internal flux. The objective vector ***γ***_*i*_ indicates which reactions within the cell contribute directly to cellular biomass, and so is non-zero only in elements corresponding to internal fluxes. We can therefore rewrite this vector to include only elements corresponding to internal fluxes, so that the objective of the optimization is to maximize ***γ***_*i*_ · ***ϕ***_*i*_.

The stoichiometry of metabolite exchange reactions is represented by standard basis vectors [9]. Therefore, we can partition Γ_*i*_ as

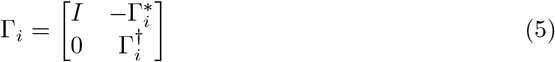

where *I* is the identity matrix of appropriate size, and 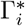 and 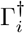 contain the stoichiometry of the internal reactions [9, 39, 40].

We can see from Eq. (5) that ker(Γ_*i*_) is isomorphic to ker 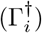, and so we can maximize over this kernel. Then, the exchange reaction fluxes are determined by the internal fluxes according to the linear mapping 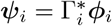. The maximization of FBA becomes a maximization problem over the internal fluxes^1^. We rewrite Eq. (4) using Eq. (5) and combine with Eqs. (2) and (3) to form the differential algebraic system

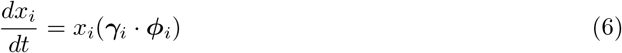

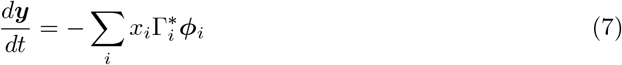

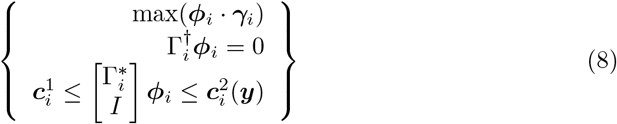

where each ***ϕ***_*i*_ is determined by the optimization Eq. (8), all carried out separately. Note that this is a metabolite mediated model of community growth as defined in [27]. That is, the coupling of the growth of the separate microbes is due to the shared pool of metabolites ***y***. Each separate optimization which determines ***ϕ***_*i*_ at a single time-step depends on ***y***, and each ***ϕ***_*i*_ determines some change in ***y***.

We write, for full generality, upper and lower dynamic bounds on internal and exchange reactions, and assume that each function *c*_*ij*_(***y***) ∈ *C*^∞^. We let

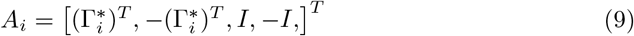

so that we can rewrite the optimization problem Eq. (8) as

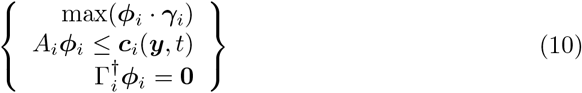

for ease of notation.

We now hope to select a basic index set 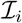 for Eq. (10) for each organism *x*_*i*_ so that each ***ϕ***_*i*_(*t*) is a solution to the resulting linear system of equations.

## Methods

### Linear optimization preliminaries

In this manuscript, we will rewrite the FBA optimization problem in the form

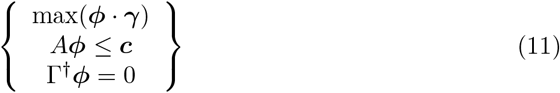

where the matrices *A* and Γ^†^ are derived from the stoichiometric matrix and flux constraints. Such a problem is often referred to as a *linear program* (LP). We now recall some well known results from the study of linear programming (see, for example [34, 41]).

First, we note that Eq. (11) can be rewritten in the so-called *standard form* with the addition of *slack variables* ***s*** = (*s*_1_, …, *s*_*n*_) which represent the distance each of the *n* constraints is from its bound as follows:

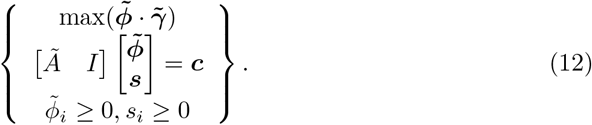

Standard form requires that we rewrite 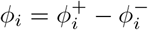 and then define 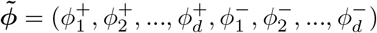 so that we require non-negativity of each variable, and the matrix *Ã* = [*A B*], *B* = −*A*. We rewrite the problem in this form to make use of established results, and for ease of notation will write ***ϕ*** instead of 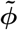 when it is clear which form of the problem we are discussing.

We will make use of the well-known result that there exists an *optimal basis* or *basic set* for a bounded linear program [42]. To state this result, we first define the notation 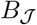 to be the matrix with columns of [*Ã I*] corresponding to some index set 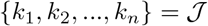, and if 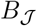 is invertible we define the notation 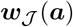 so that

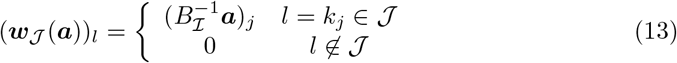

for any ***a*** ∈ ℝ^*n*^. We may now define an *optimal basis* and *optimal basic set*.

#### Definition 1.

*A* basic optimal solution *to a linear program is an optimal solution along with some index set* 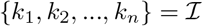 *such that* 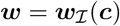, *where **c** is the vector of constraints as in* *Eq.* (12). *The variables* 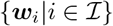 *are referred to as* basic variables, *and the index set* 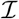 *is referred to as the* basic index set.

Finally, if there exists a bounded, optimal solution to Eq. (12), then there exists a basic optimal solution and corresponding basic index set.

For a given basic optimal solution vector ***w***, there may be more than one basic index set 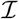 such that 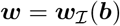. Such a solution is called *degenerate*. Clearly a necessary condition for such non-uniqueness is that there exists some 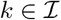 such that *w*_*k*_ = 0. This is also a sufficient condition as long as there is some column of [*Ã I*] which is not in the column space of 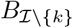.

### Forward simulation without re-solving

Consider again Eq. (10), the linear program that must be solved at each time point of the dynamical system for each microbial population. Information from prior solutions can inform future time-steps as long as the region of feasible solutions has not qualitatively changed. Thus, we may only need to solve the optimization problem a few times over the course of a simulation, rather than at every time step. The key observation making this possible is that the simplex method of solving a linear program provides an optimal basis for the solution. We may often re-use this basis within some time interval, and so find optimal solutions without re-solving the linear program.

We would like to find a form of the solution which may be evolved in time. To do this, we turn the system of linear inequalities given in the linear program into a system of linear equations. Then, if this system has a unique solution we have reduced the task to solving a system of equations rather than optimizing over a system of inequalities. We can find such a system of equations by solving the linear program once, and using this solution to create a system of equations whose solution provides the optimal flux ***ϕ***_*i*_, as described above. We then use this same system to simulate forward without the need to re-solve the LP until our solution becomes infeasible.

The linear program Eq. (10) is transformed into standard form (Eq. (12)). Then, a basic optimal solution is found with corresponding basic index set 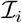. The dynamical system Eqs. (6), (7) and (10) can then be evolved in time using Eq. (13). This evolution is accurate until some *w*_*ij*_ becomes negative. Then, a new basis must be chosen. That is, until 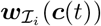 becomes infeasible, we let 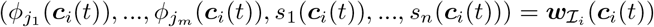 and replace Eqs. (6), (7) and (10) with

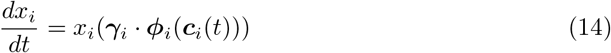

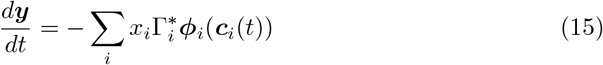

One major difficulty in this technique is that a unique ***w***_*i*_ does not guarantee a unique basis set 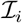. If we have some 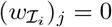 for 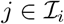, then there exists some alternate set 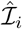 such that 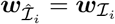. Such a solution 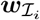 is called *degenerate*. In a static implementation of a linear program, the choice of basis of a degenerate solution is not important, as one is interested in the optimal vector and optimal value. However, as we will demonstrate with Example 1, the choice of basis of a degenerate solution is important in a dynamic problem. In fact, if the system given in Eqs. (14) and (15) is evolved forward until 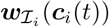 becomes infeasible, the time at which the system becomes infeasible is the time at which we have some 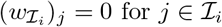. Thus, we need to resolve Eq. (10) whenever 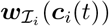 becomes degenerate, which will be the final time-point at which the 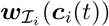 is feasible.

#### Example 1.

*Consider the dynamic linear program*

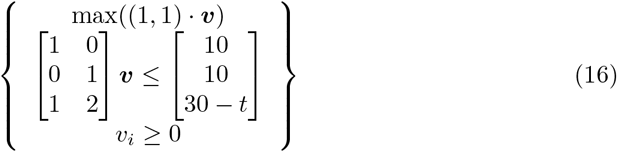

*In standard form at t* = 0, *this linear program becomes*

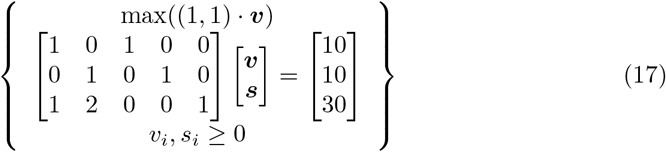

*which has the unique solution **w*** = (10, 10, 0, 0, 0). *There are three choices of basic index sets:* 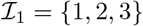, 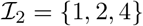, *and* 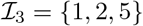. *The resulting bases are*

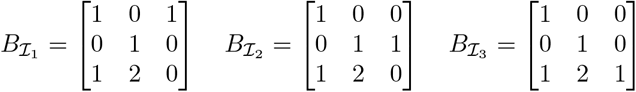

*Computing* *Eq.* (13) *at t* > 0 *for each, we have that* 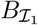 *yields* 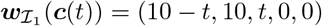, 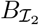 *yields* 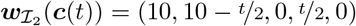, *and* 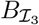 *yields* 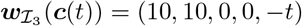, *shown in Fig. 2 for t* > 0. *Thus, only* 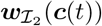 *solves the dynamic problem because* 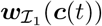 *is not optimal and* 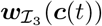 *is not feasible for t* > 0. *Notice that the correct choice of basis fundamentally depends on the time-varying bound function **c***(*t*) = (10, 10, 30 − *t*). *To see this, consider other possible time-varying bounds* ***c***(*t*) *which have **c***(0) = (10, 10, 30). *For example, if **c***(*t*) = (10 − *t*, 10 − *t*, 30), *then only* 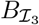 *would give the correct **w***(***c***(*t*)) *for t* > 0.

**Fig 2.**
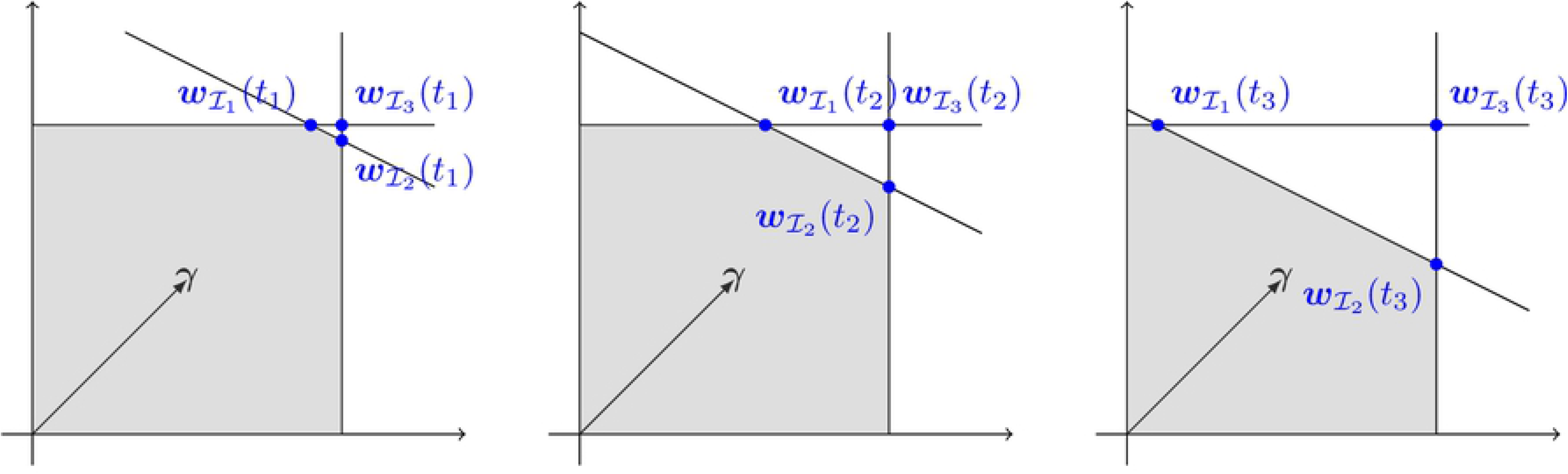
Geometric representation of Example 1 for *t*_3_ > *t*_2_ > *t*_1_ > 0, showing the three options for bases which are equivalent at *t* = 0. Note that the best choice depends on the function ***c***(*t*) = (10, 10, 30 − *t*) and cannot be chosen using the static problem alone. The feasible region of the optimization problem is shown in gray.

### A basis for the flux vector

We now provide a method to choose a basis 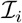 for each organism *x_i_* in the case of a degenerate solution. Consider an optimal solution ***w***_*i*_ to the linear program Eq. (12). To simulate forward according to Eqs. (14) and (15), we need for each organism *x_i_* a basic index set 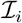 such that

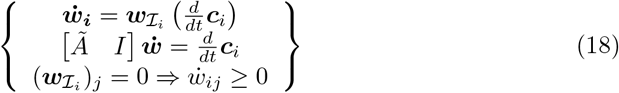

so that the solution remains feasible, and furthermore that 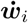 is optimal over the possible choice of basic index sets for ***w***_*i*_. This is obviously a necessary condition forward simulation within some non-empty time interval, and can be made sufficient (although no longer necessary) by making the inequality 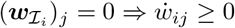 strict. We use the relaxed condition for more practical applicability.

First, we show that such a solution exists.

#### Lemma 1.

*For a linear program with the form given in* *Eq.* (12) *with a basic optimal solution **w**, there exists a basic index set* 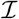 *such that* Eq. (18) *holds and* 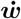 *is optimal over the possible choice of basic index sets for **w***.

*Proof*. For convenience, we now restate Eq. (12):

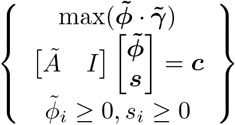

where we write 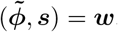.

We note that there is a finite number of basic index sets for ***w***, and so we need only show that there exists 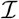 such that Eq. (18) holds. Then, the existence of an optimal such 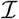 follows trivially.

If ***w*** is not degenerate, then the unique choice of basic index set 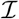 satisfies Eq. (18). To see this, simply note that if ***w*** is non-degenerate, then for every 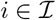, *w_i_* > 0. Thus, Eq. (18) only includes non-negativity constraints on 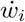 if 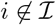, and for any 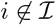, 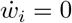. Thus, the non-negativity constraints are enforced. The equality constraints are enforced by the definition of 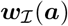 given in Eq. (13), which implies that 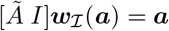 for any vector ***a*** ∈ ℝ^*n*^.

In the case of a degenerate solution ***w***, we use the following procedure to choose a set of basic variables. Let 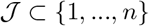 be the indices of the *n*_1_ slack variables such that *s_j_* = 0 if 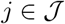 (recalling that each *s_i_* is a component of the vector ***w***). Then, let 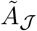 be the matrix with rows *m_j_* of 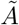 for 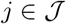. Next, let 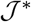 be the indices of the *n*_2_ non-slack variables such that *ϕ_j_* = 0 and 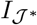 the corresponding rows of the identity matrix *I*. Notice that we now have that

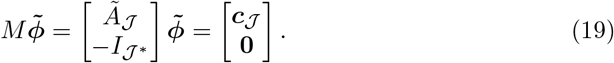

and that if *w_j_* = 0 then either 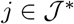 or *w_j_* = *s_k_* where 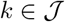 so that 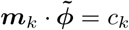 (i.e. *s_k_* is a slack variable and *s_k_* = 0). Notice that because Eq. (12) has a bounded solution, then we can assume without loss of generality that if *M* ∈ ℝ^*q*×*r*^, then *rank* (*M*) = *r* (i.e. *M* is full rank) because ***w*** must satisfy at least *r* linearly independent constraints. If this is not the case, then the problem can be projected onto a lower dimensional subspace.

Consider the linear program

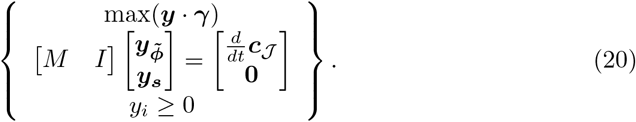

Assume that there is some basic optimal solution to Eq. (20) with a basic index set 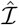 such that exactly *r* slack variables are non-basic, where again *r* = |***ϕ***| is the rank of the matrix *M*. This implies that there are *r* linearly independent rows of *M* (which we index by 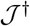) which form an invertible matrix 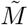 such that

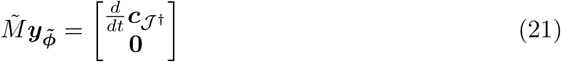

and we can then determine ***y***_***s***_ by

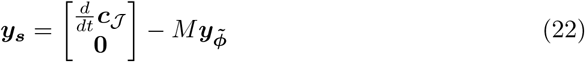

and note that each (***y***_***s***_)_*i*_ ≥ 0. We now rewrite 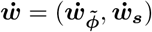 from Eq. (18) and define 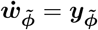 and

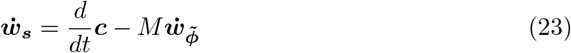

and conclude that this satisfies the constraints of Eq. (18). Next, we take 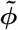 to be the unique solution to

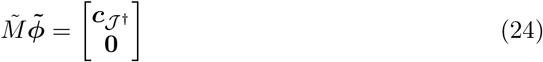

and 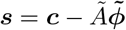.

Finally, we take 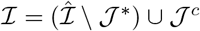 and note that this basis set enforces exactly the same *r* linearly independent constraints as 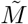^2^.

We now prove that there is some basic optimal solution to Eq. (20) with a basic index set 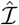 such that exactly *r* slack variables are non-basic, where *r* is the rank of the matrix *M*.

First we note that for any basic optimal solution, if there are *r*\* > *r* slack variables which are non-basic, then there are *r*\* rows of 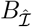 which are non-zero only in the columns of *M*. Therefore, 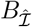 is not invertible. We can conclude that the number of non-basic slack variables is at most *r*.

Next, suppose 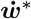 is a basic optimal solution with basis 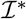 such that there are *r*\* < *r* slack variables which are non-basic.

We would like to assume that there are at least *r* slack variables 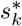 corresponding to *r* linearly independent constraints such that 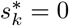. Recall that 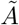 was formed with repeated (negated) columns in order write the problem in standard form (the non-negativity bounds of Eq. (12) are artificial). Therefore, we can find some vector ***x*** in the kernel of the matrix formed by the rows of 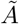 corresponding to zero slacks which also has ***x*** · ***γ*** = 0. We can therefore find a vector ***y*** in the kernel of

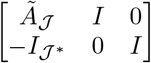

which has *y_k_* = 0 if *s_k_* = 0 and *y_j_* ≠ 0 if *s_j_* ≠ 0 and *s_j_* corresponds to a constraint that is not a linear combination of the constraints corresponding to the *s_k_* = 0. There is at least one such constraint as long as the 0 slack variables correspond to constraints with span less than dimension *r*, and so we can take 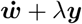 for some *λ* and so increase the number of non-zero slack variables. We can therefore assume without loss of generality that there are at least *r* slack variables 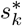 corresponding to *r* linearly independent constraints such that 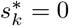, as was desired.

We can finally choose some linearly independent set of *r* constraints which correspond to 0 slack variables, and call the matrix whose rows are these constraint vectors *M*\*. Now, because there are *r*\* < *r* non-slack basic variables, there is some non-slack, non-basic variable *v_j_* such that the column 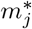 of *M*\* (and *m_j_* of *M*) is linearly independent from the columns corresponding to the *r*\* non-slack basic variables. We can conclude that if

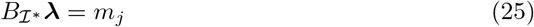

then there is some *λ_k_* ≠ 0 where *k* corresponds to the index of a slack variable with *s_k_* = 0. We can remove *k* from the basic index set and add *j* without changing 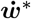, and therefore preserving optimality and feasibility. We have then increased the number of non-basic slack variables, and we can repeat if necessary to form 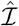 with exactly *r* non-basic slack variables.

### Implementation

We implement dynamic FBA for python in the package `surfinFBA`. The main algorithm begins by computing an in initial optimal internal flux vector each species according to Eq. (10). A secondary optimization is then carried out, which can be specified by the user. If no secondary objective is provided, the program default is to minimize total internal flux. If the optimal solution is not degenerate, we may begin simulating forward until the solution becomes infeasible, and then repeat. Either the initial solution to Eq. (10) or the solution found when the initial solution becomes infeasible will be degenerate. Then, a new basis must be chosen as implied by Lemma 1. Note that in practice, the way that we form the matrix *A* guarantees that *rank* (*A*) = *d*, where 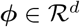 (i.e. *d* is the number of internal fluxes).

To choose among possible bases of a degenerate solution, we form a second linear program analogous to Eq. (18) in the following way. We first find all constraints ***a***_*j*_ (i.e. rows of *A_i_* or 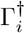) such that ***a***_*ij*_ · ***ϕ***_*i*_ = *c_ij_*(*t*), calling this set 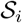. Note that this set contains all the rows of 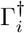, for which we regard *c_ij_*(*t*) = 0 for all *t* > 0. Note that if the solution given is a basic optimal solution, the rank of the matrix whose rows are ***a***_*ij*_ for 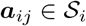 is *d*, where again *d* is the number of internal fluxes. This is true because we include constraints of the type *a* < *ϕ*_*ij*_ < *b* as rows of *A_i_*.

Then, we solve the linear program

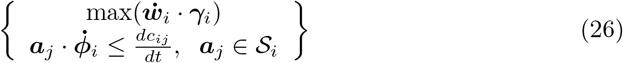

We may then use any basis 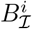 which solves Eq. (26) as long as it has exactly *d* non-basic slack variables. Lemma 1 tells us that such a choice exists, although it may be necessary to manually pivot non-slack variables into the basis set given by the numerical solver ^3^. Note that we do not need the entire basis 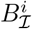, but instead only need the *d* × *d* submatrix formed by rows of *A_i_* or 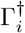 which correspond to non-basic slack variables in the solution to Eq. (26). These appear as rows ***a***_*i*_, **0**) in 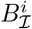, and so this sub-matrix uniquely determines ***ϕ***_*i*_. We call this smaller matrix *B_i_*, and label the set of row indices as 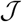.

The chosen basis 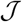 and corresponding constraints is used to simulate forward until the solution becomes infeasible. At that time, we have an optimal solution to Eq. (10) simply by continuity. We therefore do not need to resolve Eq. (10) but instead re-form and solve Eq. (26).

## Results

### Error estimation

Figure 3 implies that a simulation of a microbial community can be divided into time intervals on which the algorithm is exact. That is, there exits some sequence *t*_0_ = 0 < *t*_1_ < ··· < *t*_*n*−1_ < *t_n_* = *T* such that if we know the optimal flux vectors ***w***_*i*_(*t_l_*) at time *t_l_*, then Lemma 1 implies the existence of a set of invertible matrices 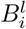 such that solutions to Eqs. (14) and (15) are solutions to Eqs. (6), (7) and (10) for *t* ∈ [*t*_*l*_, *t*_*l*+1_]. Therefore, if we are able to identify the *t_l_* exactly, then Fig. 3 provides exact solutions to the dynamic FBA problem Eqs. (6), (7) and (10). Of course, numerical limitations imply that we will not re-optimize precisely at each *t_l_*, and so we must investigate the impact of this error. However, once re-optimization is done, the method is again exact. The result is that we have no local truncation error for any time step taken between re-optimization after *t_l_* and the interval endpoint *t*_*l*+1_, except for error due to numerical integration.

**Fig 3.**
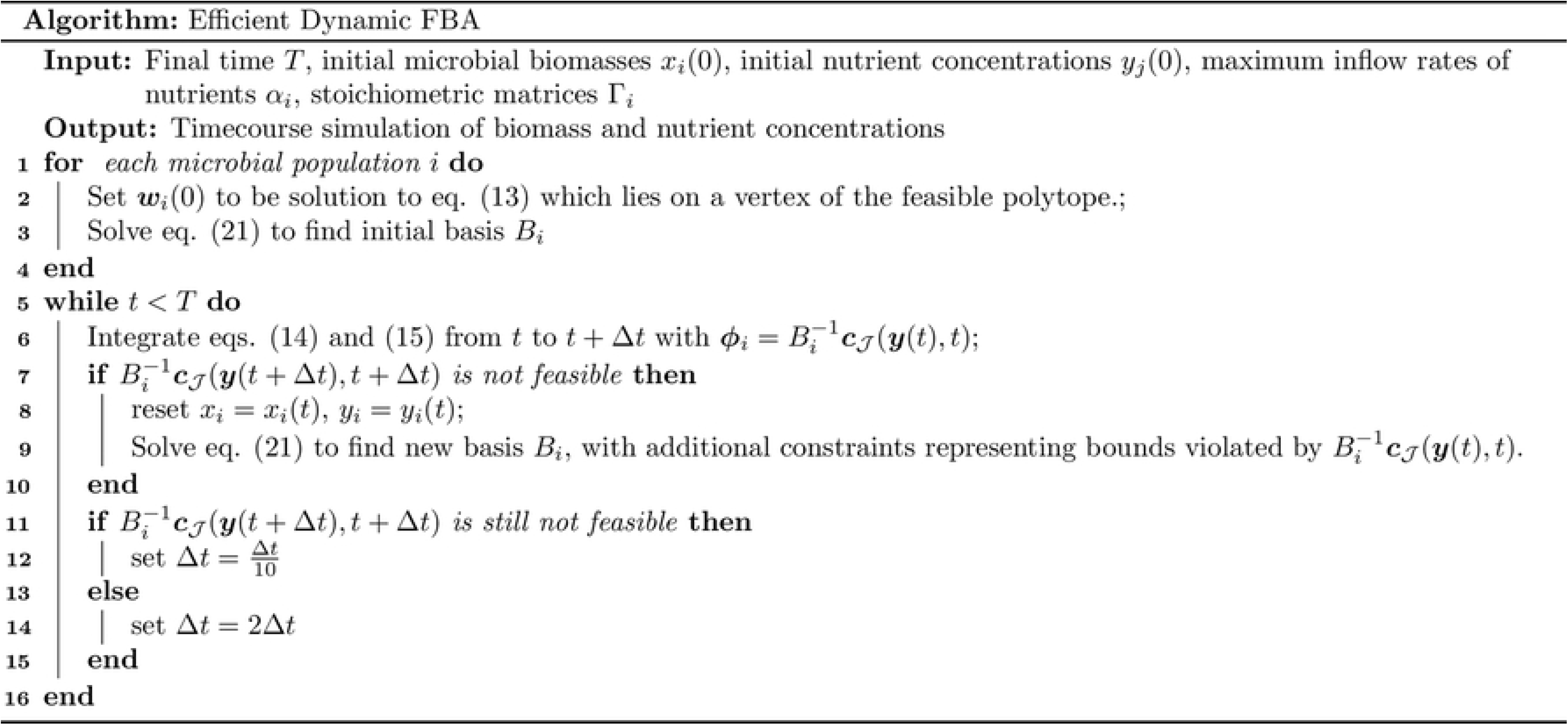
Dynamic FBA algorithm following Lemma 1. Note that for numerical stability and speed, we may store the matrices *Q*_*i*_, *R*_*i*_ such that *Q*_*i*_*R*_*i*_ = *B*_*i*_ is the QR-factorization of *B*_*i*_ rather than either storing 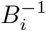 or solving completely during each time step of numerical integration.

Assume that *t*_*l*−1_ is known exactly, and *N* is such that *t*^1^ = *t*_*l*−1_ + (*N* − 1)Δ*t* ≤ *t*_*l*_ < *t*_*l*−1_ + *N* Δ*t* = *t*^2^, so that there is some possible error in the interval [*t*^1^*, t*^2^]. We can estimate the accumulated error in this time interval using a power series expansion. Let ***x***(*t*), ***y***(*t*) be solutions to Eqs. (6), (7) and (10) and 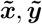 be solutions given by Fig. 3 for *t* ∈ [*t*^1^, *t*^2^). Furthermore, let 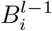 be the invertible matrices derived by solving Eq. (10) at *t*_*l*−1_ and 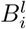 those derived by solving at *t_l_*. Then, 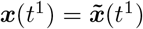 and 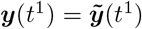. For each *x_i_* we expand, assuming some regularity of the functions ***c***(***y***)

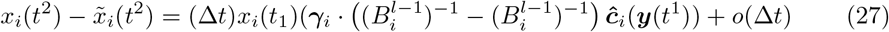

and see that this method gives first order local error in time steps that require a re-optimization.

The local error, while first order, only appears at time steps in which a re-optimization occurred, and so global error will scale with the number of necessary re-optimizations. This is in contrast with Fig. 1, which gives first order local error at every time-step.

### Number of optimizations

One way in which we can assess the usefulness of Fig. 3 in relation to the classical algorithm (Fig. 1) is to estimate the number of times a linear program must be solved in a simulation with a given error tolerance. In addition to the obvious relationship with the speed of the simulation, the number of re-optimizations needed also effects the accuracy of Fig. 3, as we showed in the previous section.

Clearly, Fig. 1 requires *N* linear programs to be solved, where *N* is the number of time steps taken in the simulation. On the other hand, we see in Fig. 4 that Fig. 3 requires a linear program be solved far fewer times. See supplemental `S1_File.pdf` for details on community simulations using both Fig. 3 and Fig. 1.

**Fig 4.**
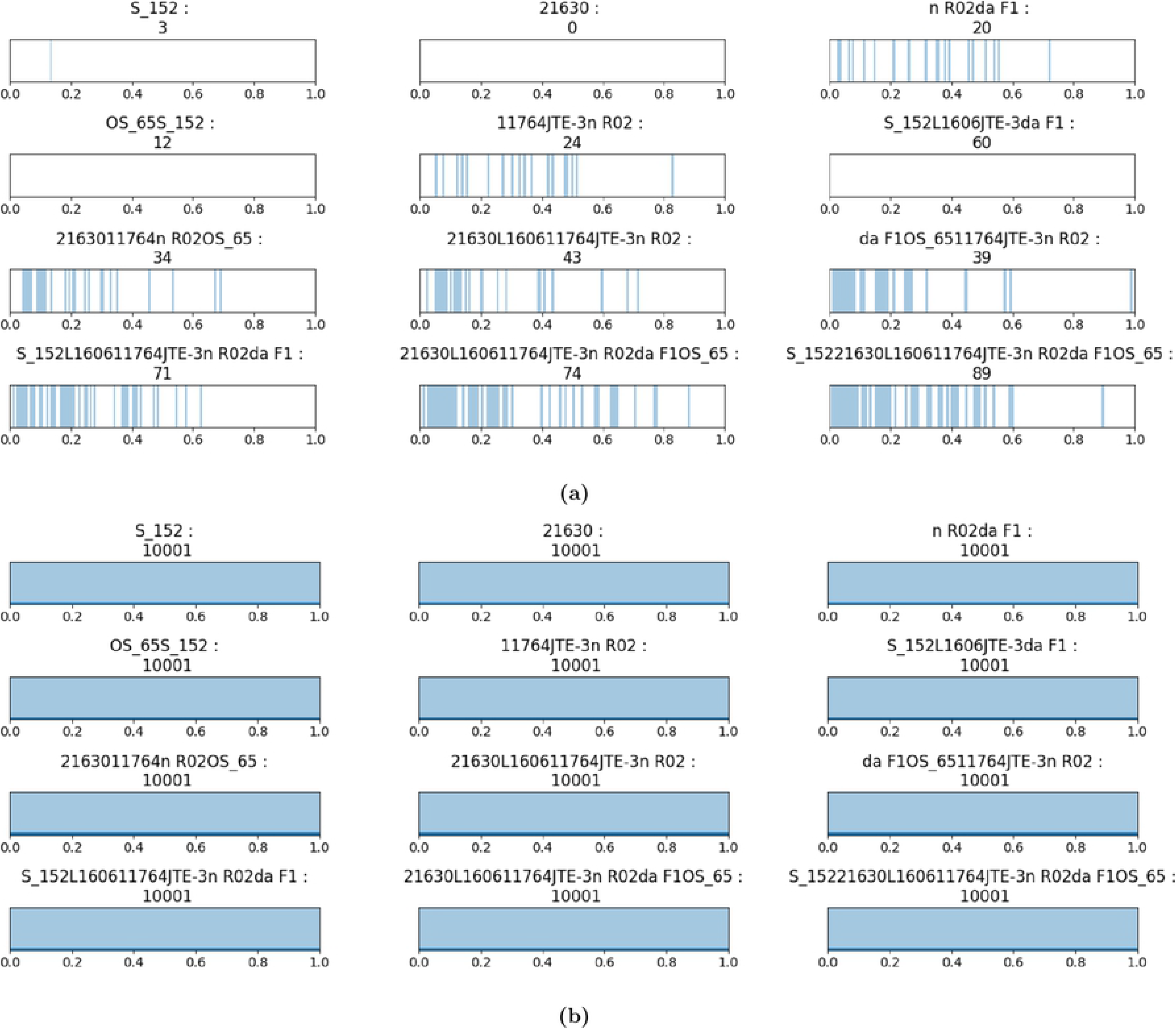
(a) Time-points of re-optimizations required in simulations using Fig. 3. (b) Time-points of re-optimizations required in simulations using Fig. 1. For model. See supplemental file S1 for models used and wall time of computation. The number of re-optimizations is given in each plot title.

### Computational time

We can also compare implementations of Fig. 3 and Fig. 1. Figure 1 is analogous to Euler’s method for computing solutions to ordinary differential equations, and is likewise depended on time-step size for both speed and accuracy. Simulating communities of more than a single organism required a time step size of 10^−4^ in order to avoid numerical error which caused the solution to terminate. For too large a step size, numerical error caused metabolite concentrations to become less than zero and the simulation to terminate. To compare Fig. 3 and Fig. 1, we implemented Fig. 1 for communities using the FBA methods of the COBRAPy Python package [8]. The exact implementations of Fig. 3 and Fig. 1 that we compared are available for download as a Python package at https://github.com/jdbrunner/surfin_fba.

Figure 5 (a) shows the wall time to simulate dynamic flux balance analysis with no microbial dilution, metabolite dilution, or metabolite replenishing for various small communities. To test the algorithms, we assumed linear nutrient uptake with randomly drawn nutrient uptake rates. That is, in Eq. (10) we took for each organism *i*

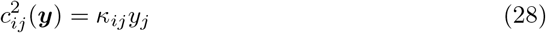

with *κ_ij_* ∈ (0, 1) chosen uniformly at random. For each community and choice of {*κ_ij_*}, we simulated using both algorithms and recorded wall time to simulate. Figure 5 (b) shows the time length of the simulation per organism. We can see that, for small communities at least, the time to simulate using Fig. 3 scales linearly with community size.

**Fig 5.**
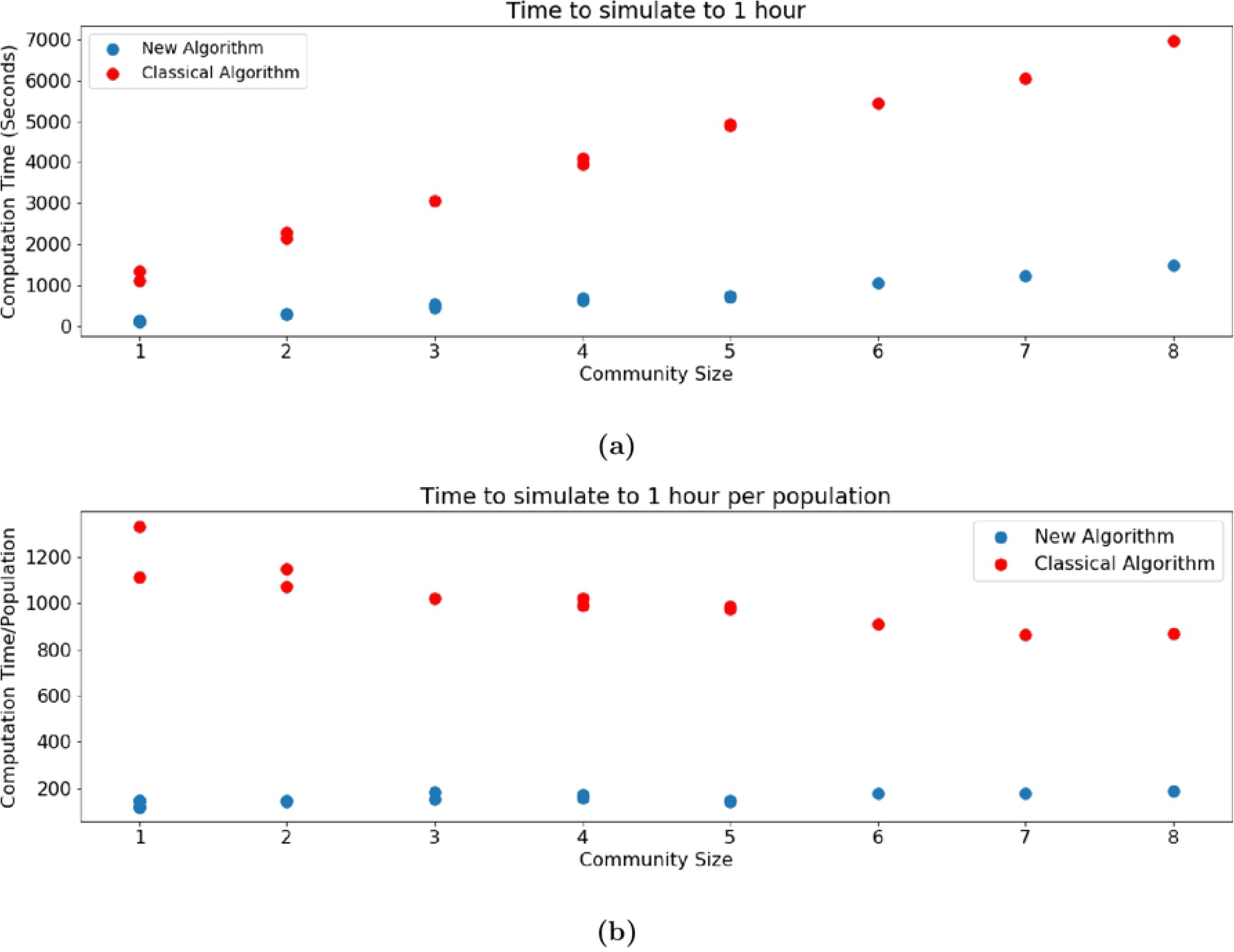
(a) Comparison of time to simulate community models to *t* = 1, with no inflow or outflow of metabolites or microbes. (b) Comparison of time to simulation per community size of community models with no inflow or outflow of metabolites or microbes.

## Examples & applications

### An 8 species community model

Our method can be used to simulate a community of eight microorganisms using “off the shelf” models downloaded from the online bacterial bioinformatics resource *PATRIC* [18]. We simulate the eight species used in Friedman et. al [21] using Fig. 3 in a low flow chemostat with random metabolite uptake rates and random microbial death rates. The community dynamics are shown in Fig. 6.

**Fig 6.**
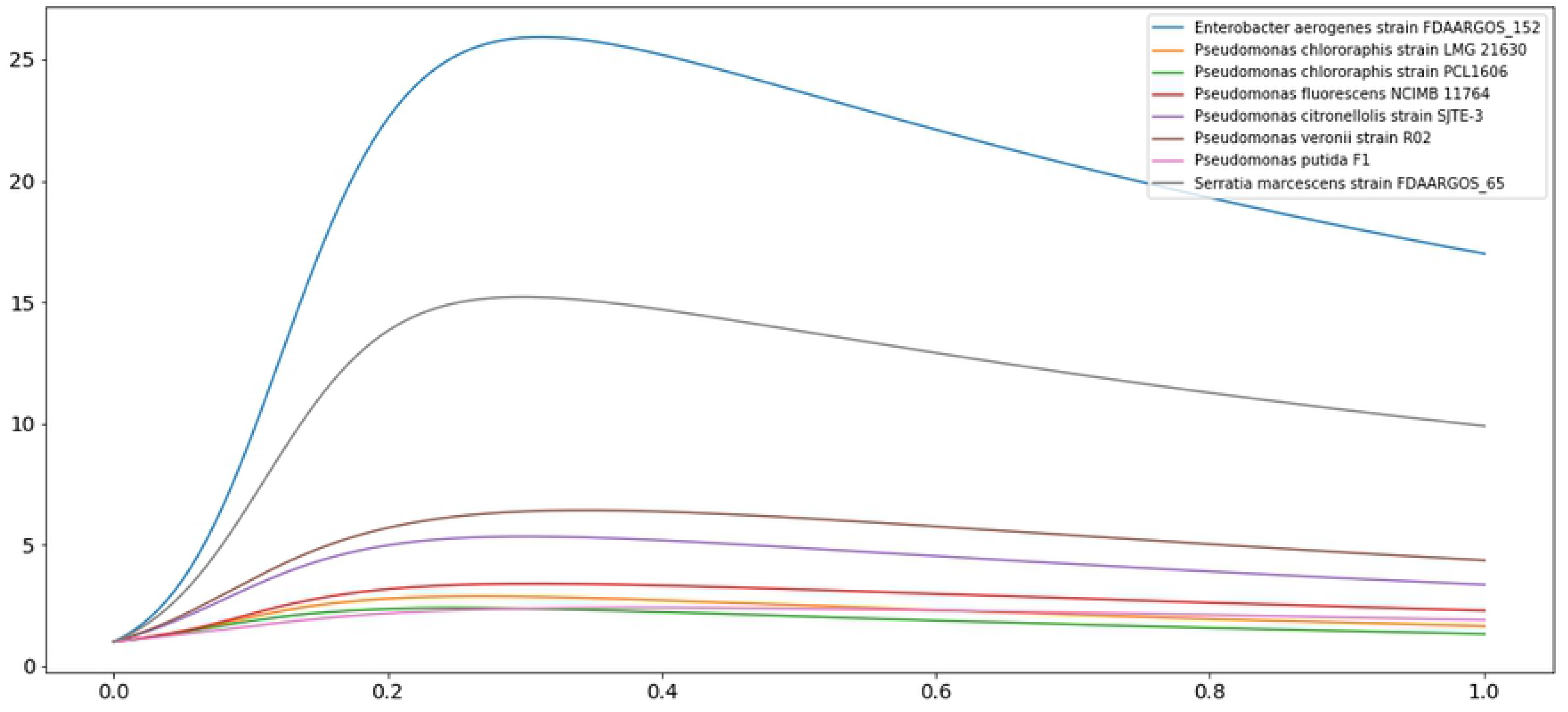
8 Species low-flow chemostat community model with random microbial death rates and metabolite uptake rates. This is model *S 15221630L160611764JTE-3n R02da F1OS 65*, which simulates the a community of the 8 species used in Friedman et. al [21] using Fig. 3. Metabolic models for each species were downloaded from the online bacterial bioinformatics resource *PATRIC* [18].

We can also inspect the dynamics of the metabolite biomass in the medium. Interestingly, many metabolites show non-monotonic dynamics, and thus interesting transient behavior before equilibrium is reached. In Fig. 7, we show one such metabolite, uracil. Furthermore, we can inspect the effect microorganism *i* has on the metabolite over time by inspecting 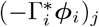, where *j* corresponds to this metabolite. For example, we see that *P. citronellolis* initially consumes uracil 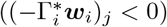, but later switches to producing the metabolite.

**Fig 7.**
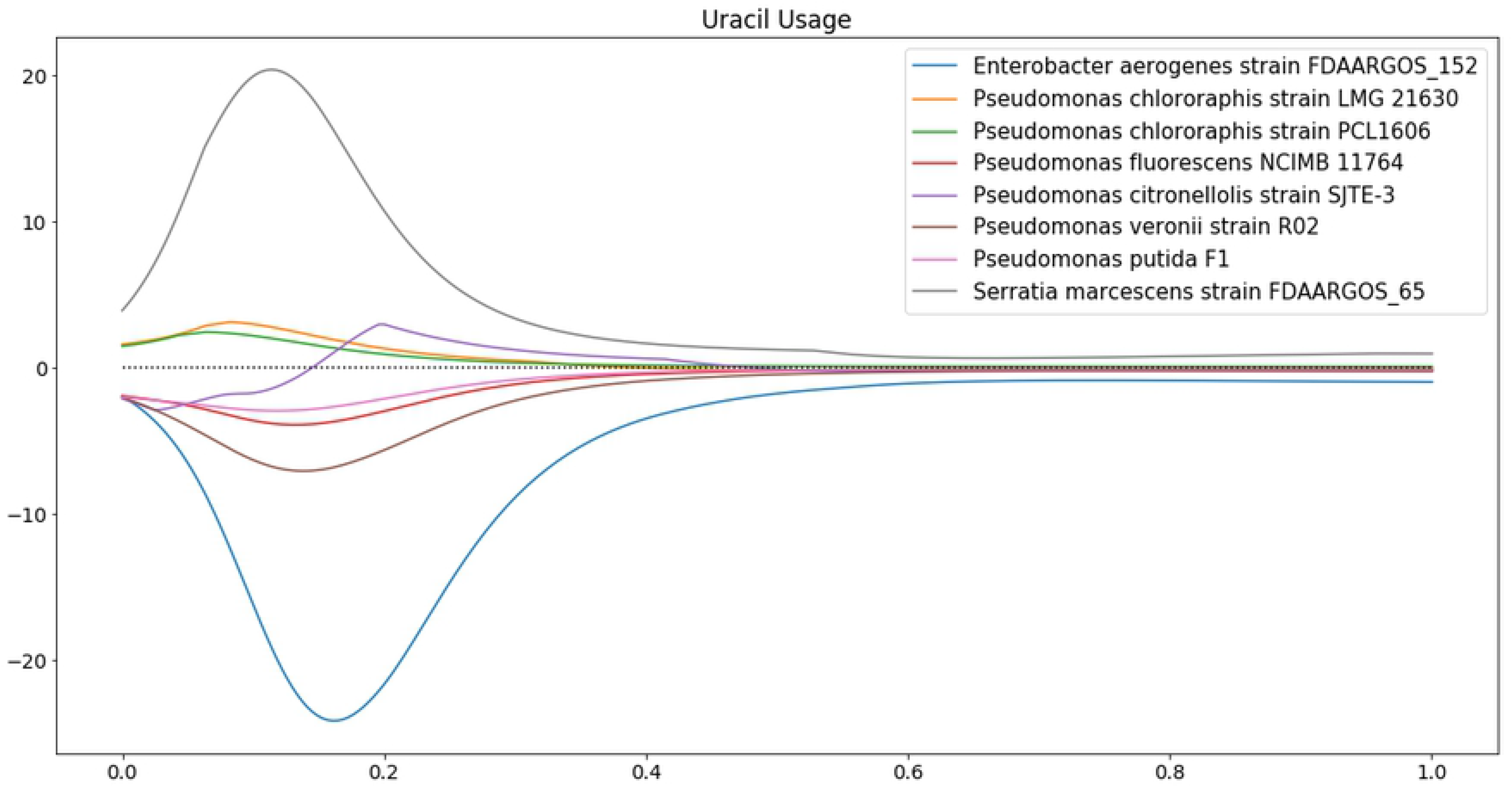
In the eight species community model shown in Fig. 6, uracil biomass changes in complicated ways. We can see which microorganisms are having an effect on this metabolite by inspecting each organism’s contribution to 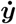. When this is positive, this organism is producing the metabolite, and when it is negative, the organism is consuming the metabolite. Interestingly, *P. citronellolis* transitions from initially consuming to later producing uracil.

### Prediction dependence on nutrient uptake

It is worth noting that dynamic FBA depends fundamentally on the nutrient uptake bounds ***c***(***y***) in Eq. (10). For our simulations, we take these to be linear,

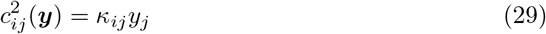

meaning that the maximum uptake rate of nutrient *y_j_* by organism *x_i_* is proportional to the concentration of *y_j_*. The choice of *κ_ij_* may have a profound effect on the outcome of a community simulation, as it represents how well an organism can sequester a resource when this will optimize the organism’s growth. In order study this effect in a small community, we sampled a three-species community model with *κ_ij_* ∈ (0, 1) chosen uniformly at random. We used models for *Pseudomonas putida F1*, *Pseudomonas chlororaphis strain PCL1606* and *Pseudomonas veronii strain R02* downloaded from the online bacterial bioinformatics resource *PATRIC* [18]. We simulated with no dilution of metabolites or microbes, and no replenishment of nutrients. In every simulation, some critical metabolite was eventually depleted and the organisms stopped growing. We recorded the simulated final biomass of each organism from each simulation, and the results are shown in Fig. 8. While a wide variety of outcomes was observed, these did cluster about a result in which *P. veronii* outgrew the other two organisms. This suggests that *P. veronii* has some competitive advantage in the COBRAPy default simulated medium.

**Fig 8.**
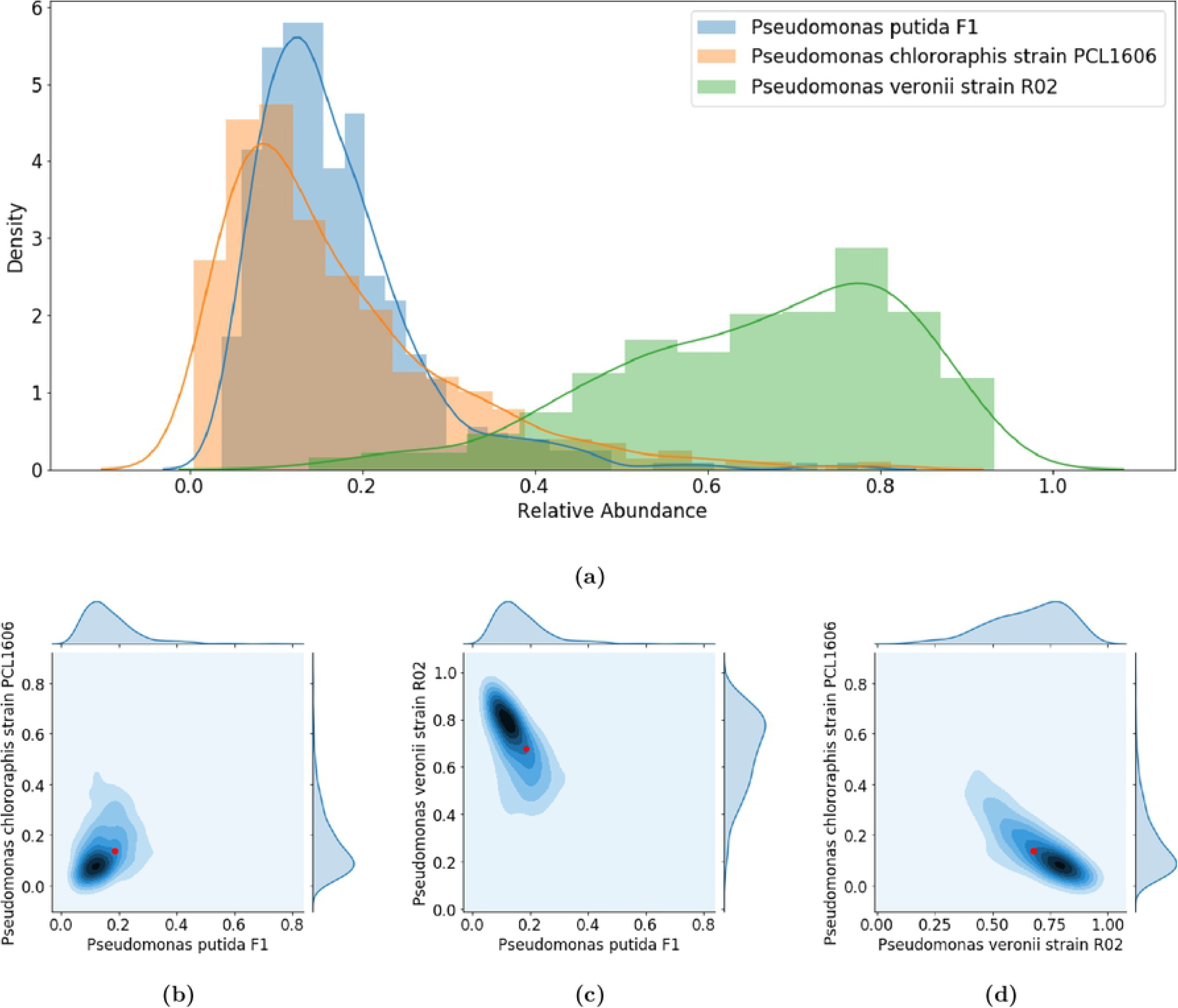
(a) Histogram of the final simulated biomass of each of *P. putida*, *P. chlororaphis* and *P. veronii* from 537 simulations, each with different metabolite uptake rates *κ*_*ij*_. (b,c,d) Pair-wise comparison of the final simulated biomass densities using a kernel density estimation. In red is the result of uniform uptake rates *κ*_*ij*_ = 0.5 for all *i, j*.

### Community growth effects

As we saw in previous section, community growth outcomes depend on the choice of nutrient uptake rates *κ_ij_*. Using Fig. 3, we can perform monte-carlo sampling in order to understand the possible effects on some microorganism of growing in some community. To do this, we randomly sample the set of uptake rates *κ_ij_* and run simulations of various communities for the chosen uptake rates. Then, the correlation between communities of final simulated biomass of some organism can be interpreted as the effect of the community on the growth of that organism. A correlation less than 1 between growth of an organism in different communities indicates that the community is having some effect. To see the direction of this effect, we can fit a simple linear regression model (best fit line) to the final simulated biomasses. Then, the slope of this line tells us if the organism benefits or is harmed by being in one community over another.

We again simulated *P. putida*, *P. chlororaphis* and *P. veronii* using models from *PATRIC*. Each organism grew to a larger final simulated biomass when alone compared to when in a trio with the other two. However, this difference was much less for *P. veronii*, which in general out-competed the other two species. These results are shown in Fig. 9.

**Fig 9.**
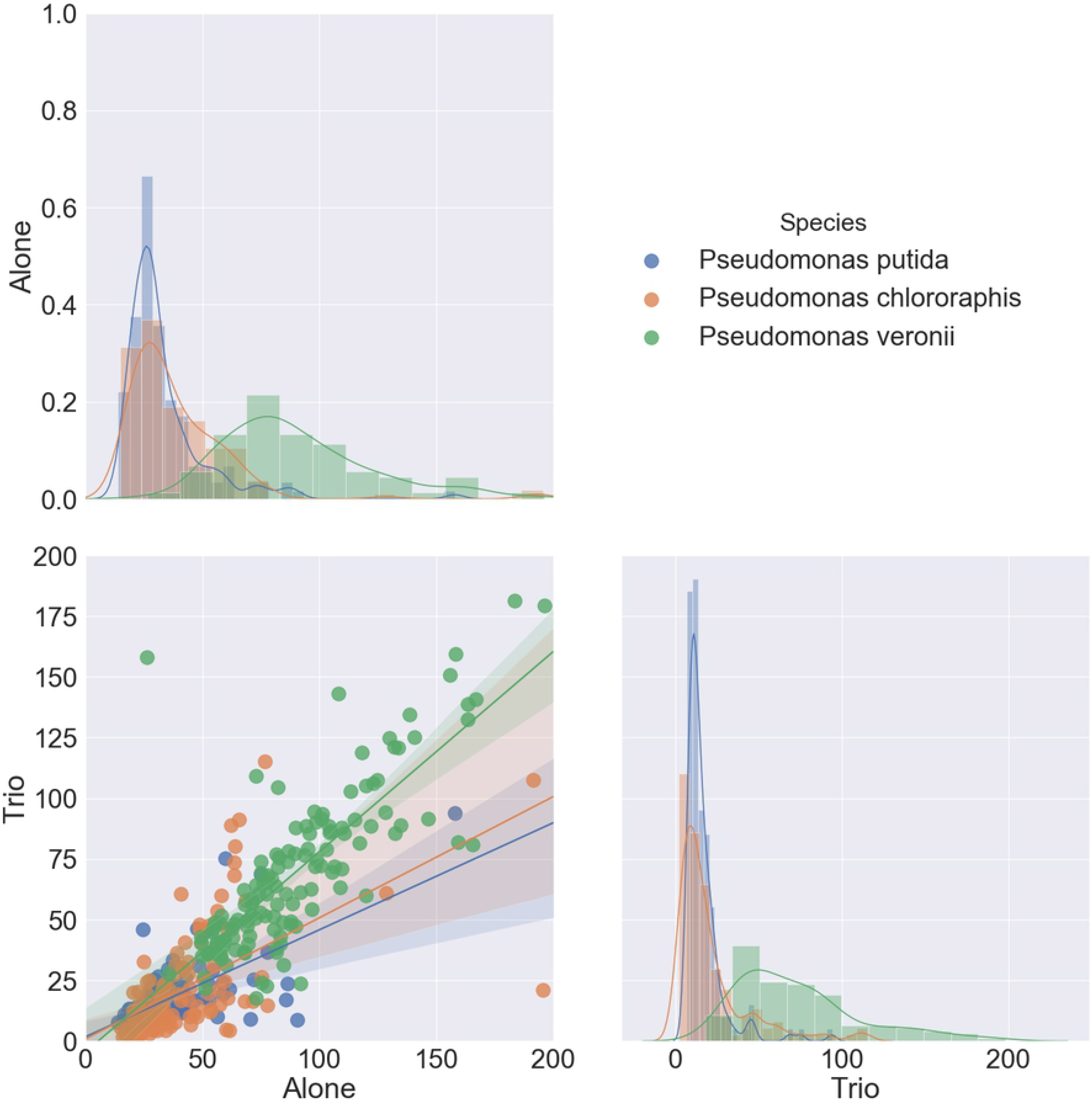
Randomly sampled uptake parameters can give an indication of metabolite mediated community effects.

## Supporting information

### S1 File. Data from test simulations

Test simulations were performed using models downloaded from *PATRIC* of species *E. aerogenes*, *P. chlororaphis*, *P. flourescens*, *P. citronellolis*, *P. veronii*, *P. putida*, and *S. marcescens*.

### S2 Software

https://github.com/jdbrunner/surfin_fba. Available code for the algorithm described in the Python language. This package is designed for use with the popular COBRAPy package for metabolic models.

## Acknowledgments

This work was supported by funding from the DeWitt and Curtiss Family Foundation, National Cancer Institute grant R01 CA179243, and the Center for Individualized Medicine, Mayo Clinic.

## Footnotes

^1^In fact, we can project onto the kernal of the matrix 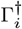 and so reduce the dimensionality of the problem. However, in practice this projection is not numerically stable.

^2^In practice, we may simply use 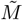 to find 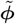

^3^In testing the algorithm, this was necessary when using IBM ILOG CPLEX Optimization Studio to solve, but not when using The Gurobi Optimizer.

